# Conformational plasticity and allosteric communication networks govern Shelterin protein TPP1 binding to human telomerase

**DOI:** 10.1101/2023.02.02.526655

**Authors:** Simone Aureli, Stefano Raniolo, Vittorio Limongelli

## Abstract

The molecular binding interaction between the Shelterin complex protein TPP1 and human telomerase enzyme (TERT) triggers the telomerase maintenance mechanism that marks cell lifespan. The TPP1’s structural element deputed to bind TERT is the OB-domain, which is able to interact with TERT’s hTEN (TPP1 binding telomerase domain) through the TEL-patch, a group of amino acids whose mutations provoke harsh pathologies. Indeed, aberrations in the formation of TPP1-TERT heterodimer can lead to severe diseases like Hoyeraal-Hreidarsson syndrome (HHS), whose patients are affected by short telomeres and extremely poor life expectancy. In the present study, we provide a thorough characterization of the structural properties of the TPP1’s OB-domain by combining data coming from microsecond-long molecular dynamics calculations, time-series analyses, and graph-based networks. Our results show that the conformational plasticity of the TPP1’s TEL-patch region is influenced by a network of long-range amino acid communications, needed for the proper TPP1-hTEN binding. Furthermore, we reveal that in the Glu169**Δ** and Lys170**Δ** TPP1 variants, responsible for HHS, the plasticity of the TEL-patch region is reduced, affecting the correct binding to hTEN and in turn the telomere processivity, which eventually leads to accelerated ageing of affected cells. Our study provides an unprecedented structural basis for the design of TPP1-targeting ligands with therapeutic potential against cancer and telomerase deficiency diseases.

## 1 Introduction

Telomeres are an ensemble of proteins, noncoding DNA, and RNA that protects chromosomes’ termini from unwanted events, like recombination and degradation, thus being crucial for cell lifespan (Fig. 1A) [1–5]. While in normal somatic cells telomeric DNA progressively shortens with each round of cell division leading to cell senescence, in cancer cells the telomeric DNA length is preserved, allowing the tumor to continue proliferating. In particular, the telomere lengthening is insured in about 85% of cancer cells by the overexpression of the telomerase enzyme [6, 7], which adds hexanucleotidic sequences to the 3’ ends of chromosomes [8]. Telomerase is a ribonucleoprotein complex consisting of a protein subunit (TERT) that works as a reverse-transcriptase using a specific RNA component (TR) as template (Fig. 1B). Four domains characterize TERT, three of which form the conserved ring-shaped catalytic core and are i) the *“Telomeric RNA Binding Domain”* (TRBD); ii) the *“Reverse Transcriptase”* (RT); and iii) the *“C-Terminal Extension*” (CTE) [9]. The fourth domain is the *“Telomerase Essential N-terminal*” (TEN) domain, deputed to enhance telomerase processivity and its recruitment to telomeres [10–13]. During its action TERT is assisted by several regulatory proteins, among these the Shelterin complex protein TPP1 plays a key role (Fig. 1C) [14–17]. In fact, TERT associates to telomeres by interacting with the *“Oligosaccharide/Oligonucleotide-binding”* domain (OB-domain) of TPP1 [6, 15, 18, 19], forming a binary complex crucial for telomere processivity. The region of the OB-domain responsible for binding to telomerase is named *“TPP1 E and L rich-patch”* (TEL-patch) due to the abundance of glutamate and leucine residues in this region (Glu168, Glu169, Glu171, Leu183, Leu212 and Glu215) [18]. Indeed, mutation or deletion of residues in the TEL-patch, or close to this region, might lead to pathologies as in the case of *“Hoyeraal–Hreidarsson syndrome*” (HHS), a severe form of dyskeratosis congenita. In such a case, the deletions of Glu169 and Lys170 (Glu169Δ and Lys170Δ, respectively) induce a reduced TERT recruitment with the consequent shortening of telomeres [20–23]. Patients affected by this disorder exhibit growth retardation, microcephaly, cerebellar hypoplasia, immune deficiency, aplastic anaemia, and bone marrow failure with poor life expectation [20, 21]. In this scenario, the understanding of the molecular basis of HHS is the first, paramount step toward the development of efficacious treatments. This requires the elucidation of the effect of Glu169Δ and Lys170Δ in TPP1 on the formation of the binary complex with hTEN. The recently disclosed cryo-EM structures of TPP1/TERT structures [24, 25] have prompted us to investigate the functional conformational changes of TPP1 in its wild-type form (WT) as well as in the Glu169Δ and Lys170Δ variants, and how these impact on the binding with hTEN. In particular, we have investigated the structural properties of TPP1 OB-domain using state-of-the-art computational techniques and the available experimental data. First, we elucidated the dynamical and structural properties of WT TPP1 by means of extensive atomistic molecular dynamics (MD) simulations. Then, MD simulations carried out upon the HHS-prone Glu169Δ and Lys170Δ variants of the OB-domain were compared with those on the WT form through time-series data analysis, including principal component analysis (PCA), cross-correlation analysis (CCA) and graph-based structure network analysis. Such investigations revealed the presence of a long-range communication network between different regions of the OB-domain and the TEL-patch. An alteration of such allosteric network leads to a decreased binding affinity of TPP1 towards TERT, with the consequent inhibition of telomere processivity. Evidence of this phenomenon is given by the Glu169Δ and Lys170Δ phenotypes - characteristic of HHS patients - that induce a large rearrangement of the TEL-patch, weakening its binding interaction with hTEN, as demonstrated by protein-protein docking calculations. An explanatory movie of the effect of Lys170Δ on TPP1 functional dynamics is available in the Supplementary Materials. The ensemble of structures and the interaction network between amino acids of WT TPP1 and its pathological variants, offers a molecular rationale and an unprecedented basis for the understanding of the functional mechanism of TPP1 and the Shelterin complex more in general. Such wealth of data paves the way to structure-based drug discovery campaigns where compounds designed to work as allosteric enhancers of the TPP1-TERT binding interaction might contrast telomeropathies like HHS, while compounds acting as allosteric inhibitors of the TPP1-TERT binary complex might represent a novel generation of anticancer agents.

**Fig. 1.**
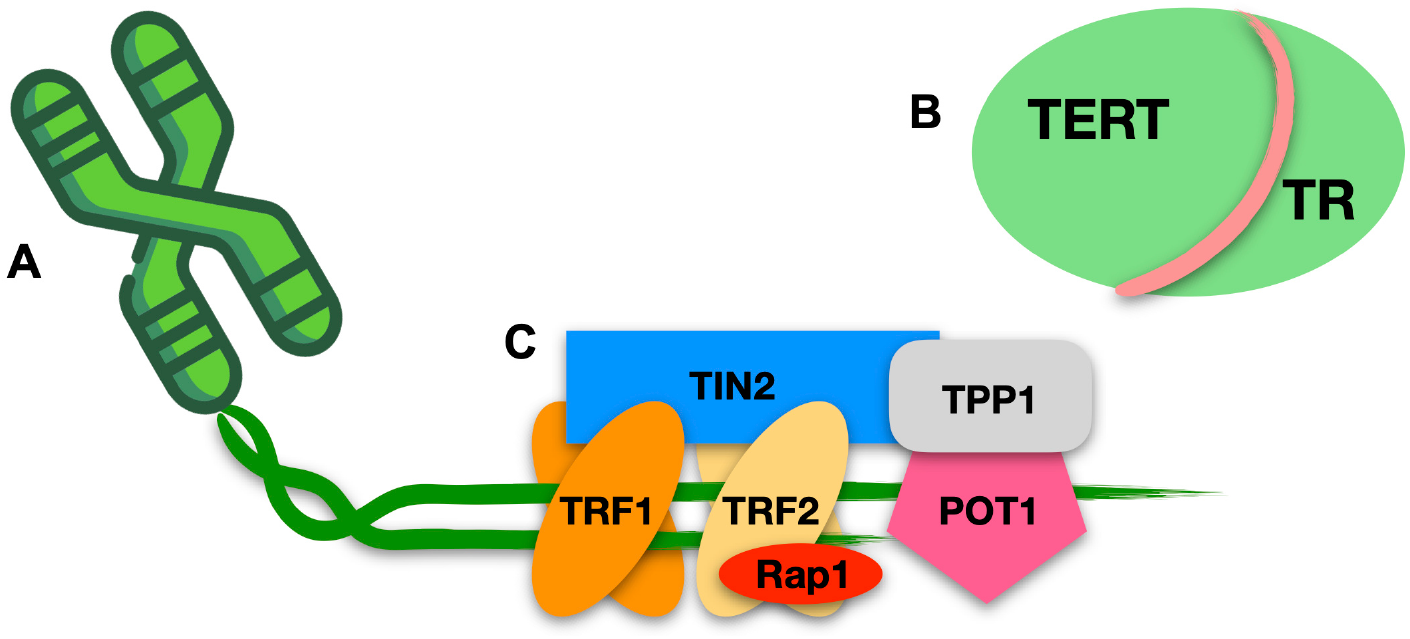
Telomeres and the telomeres’ proteome. (A) Telomeres are portions of non-coding DNA located at the chromosome termini. (B) The Telomerase ribonucleoprotein, composed of the enzyme TERT (green) and the telomeric RNA TR (pink). (C) The 6-membered Shelterin complex, composed by the double-strand binding proteins RAP1 (red), TRF2 (yellow), TRF1 (orange), and TIN2 (dodger-blue), together with the single-strand binding proteins TPP1 (grey) and POT1 (magenta).

## 2 Results and Discussions

The scope of our study is to characterize the structural properties of the shelterin protein TPP1 under physiological and pathological conditions (i.e., HHS Glu169Δ and Lys170Δ variant). To this end, we carried out a total of 3.0 *μs* atomistic MD calculations on the WT, Glu169Δ and Lys170Δ TPP1 OB-domain (Fig. 2A), and performed a number of time-series analyses on the simulated data. The results are discussed in the following paragraphs.

**Fig. 2.**
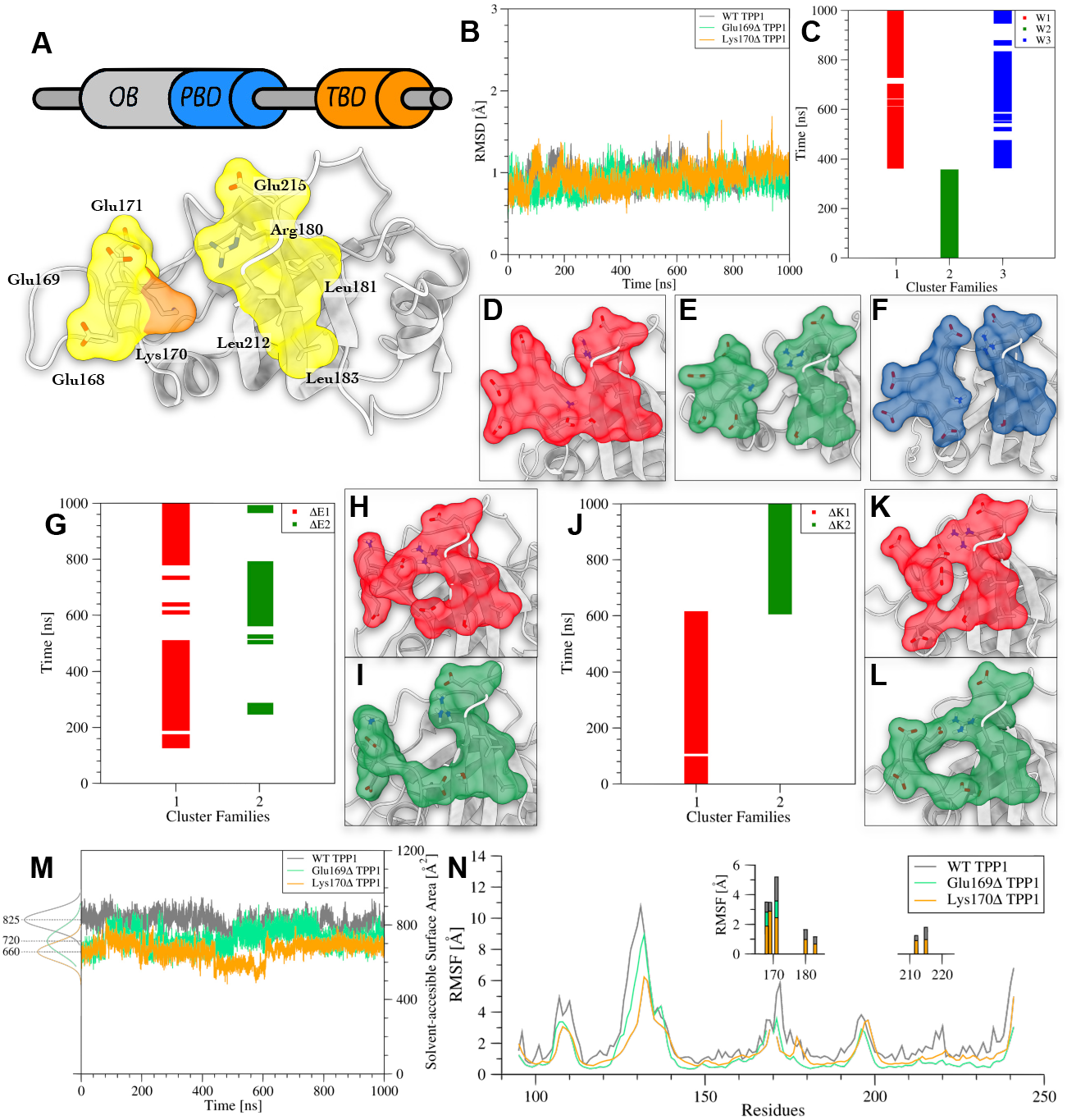
Comparison between structural features of WT TPP1, Glu169Δ TPP1, and Lys170Δ TPP1. (A) Topology of TPP1 and the 3D structure of the OB-domain (grey). TEL-patch and Lys170 are represented through their solvent accessible surface (yellow and orange, respectively). (B) Plot of the RMSD as a function of simulation time computed for the secondary structure Cα atoms of WT, Glu169Δ and Lys170Δ TPP1. (C) Exploration of WT TPP1 cluster families during the simulation. (D-F) Conformations assumed by the TEL-patch in cluster families W1, W2, and W3. Solvent accessible surface is shown in red, green and blue, respectively. (G) Exploration of Glu169Δ TPP1 cluster families during the simulation. (H, I) Conformations assumed by the TEL-patch in cluster families ΔE1 and ΔE2, respectively. Solvent accessible surface is shown in red and green. (J) Exploration of Lys170Δ TPP1 cluster families during the simulation. (K, L) Conformations assumed by the TEL-patch in cluster families ΔK1 and ΔK2, respectively. Solvent accessible surface is shown in red and green. (M) Time evolution of the SASA calculated for the TEL-patch in WT TPP1 and Lys170Δ TPP1. (N) RMSF values computed for each residue of WT, Glu169Δ, and Lys170Δ TPP1. TEL-patch’s residues are represented as histograms in the insets.

### 2.1 Structural properties of WT, Glu169Δ and Lys170Δ TPP1

The initial structures of the three TPP1 variants underwent 1.0 μs long classical MD calculations each. We first computed the Root Mean Square Deviation (RMSD) of the backbone atoms of the secondary structures during the simulations in order to inspect the overall conformational behavior of the three proteins. In Fig. 2B, the low average RMSD values (~ 1.0 Å) computed for the secondary structure Cα atoms as a function of the simulation time indicate good conformational stability for all systems. This result is further confirmed by the cluster analysis of the conformational states visited by the proteins during the MD simulations that led to a very small number of cluster families (see “Methods” section for details). However, few differences between WT, Glu169Δ and Lys170Δ TPP1 arise at the level of the TEL-patch region. At this regard, in WT TPP1 the three most populated conformation families, namely W1, W2 and W3 (Fig. 2C), present a different state of the TEL-patch. In particular, W1 shows *a closed conformation* (Fig. 2D), W2 an *open conformation* (Fig. 2E), while W3 *a semi-open conformation* (Fig. 2F). This finding reveals a certain mobility of the protein in this region that might be functional to adapt the TPP1 shape to the binding partner hTEN during the formation of the TPP1-hTEN binary complex. The structures of W1, W2, and W3 are released as PDB files in the SI. On the other hand, Glu169Δ TPP1 and Lys170Δ TPP1 show a remarkable reduction of the conformational freedom in the TEL-patch. Regarding Glu169Δ TPP1, only two conformational states were obtained from the cluster analysis, namely ΔE1 and ΔE2 (Fig. 2G). Both present the TEL-patch in the *closed conformation* (see Fig. 2H and Fig. 2I). The structural analysis of these two states during the evolution of the MD simulation reveals the presence of two long-lasting interactions formed by Glu168 with Arg180, and by Asp166 with Ser210, both absent in WT TPP1. In more detail, the Glu168-Arg180 salt bridge characterizes the ΔE1 state, while ΔE2 is stabilized by the charge-enforced H-bond formed by the side chains of Asp166 and Ser210. A similar scenario was found in Lys170Δ TPP1, for which two main conformational states were identified through the cluster analysis and named ΔK1 and ΔK2 (Fig. 2J). The analysis of their structures revealed the presence of salt bridges between the TEL-patch residues that favour the *closed conformation*. In particular, the ΔK1 state is stabilized by the Glu169-Arg180 interaction, while the salt bridge Glu171-Arg180 characterizes ΔK2 (Fig. 2K-L). Interestingly, we found out that in both variants, the deletion of a charged residue - despite opposite in charge - induces the formation of strong long-lasting interactions able to stabilize the TEL-patch in a closed conformation, thus significantly reducing the TPP1 plasticity (see Fig. S1). The structures of ΔE1, ΔE2, ΔK1 and ΔK2 are released as PDB files in the SI.

Prompted by this finding, we decided to further investigate how these differences at atomistic scale could affect the overall motion of TPP1. Therefore, we computed the solvent accessible surface area (SASA) of the TEL-patch and the root mean square fluctuation (RMSF) of each residue of WT, Glu169Δ and Lys170Δ TPP1 during the MD simulations. In line with what we previously found, both Glu169Δ TPP1 and Lys170Δ TPP1 display lower SASA values than WT TPP1 (720-660 Å^2^ vs 825 Å^2^), indicating a more compact state of the variants (Fig. 2M). The same behavior was confirmed by the RMSF calculation that shows lower values in Glu169Δ TPP1 and Lys170Δ TPP1, especially in the TEL-patch (Fig. 2N). Interestingly, our results show that the effect of the deletions is not limited to this region. In fact, both provoke long-range allosteric effects in TPP1, reducing the conformational flexibility of regions distant from TEL-patch, like that ranging from Asp123 to Gly141 (see Fig. 2N). This observation led us to perform a characterization of the TPP1 functional dynamics that is discussed in the following section.

### 2.2 Effect of Glu169Δ and Lys170Δ on TPP1 functional dynamics

In order to elucidate the effect of the Glu169 and Lys170 deletions on the functional dynamics of TPP1, we performed a principal component analysis (PCA) on the Cα atoms of TPP1 in the WT and the variant forms. Such a method allows identifying the key components (i.e., atoms) of a system that are responsible for large-scale motion endowed with a relatively long time-scale [26]. In our case, using such an approach, we could detect the most relevant slow motion of the systems along the MD simulations and finally provide a graphical representation of the main conformational rearrangements in the three variants of TPP1 (Fig. 3). Specifically, we focused our analysis on three regions of TPP1 defined as follows:

- *“Asp123-Gly1ļ1*“ (**I**), the longest loop of the TPP1 OB-domain structure (red in Fig. 3A);
- *“Thr158-Glu178*” (**II**), hosting the TEL-patch *“Knuckle*” motif, proposed by Grill et al [22], where Glu168, Glu169, Lys170 and Glu171 are located (green in Fig. 3A);
- *“Asp207-Val221*“ (**III**), where additional two residues belonging to the TEL-patch “*Barrel part*”, namely Leu212 and Glu215, are located (blue in Fig. 3A).

**Fig. 3.**
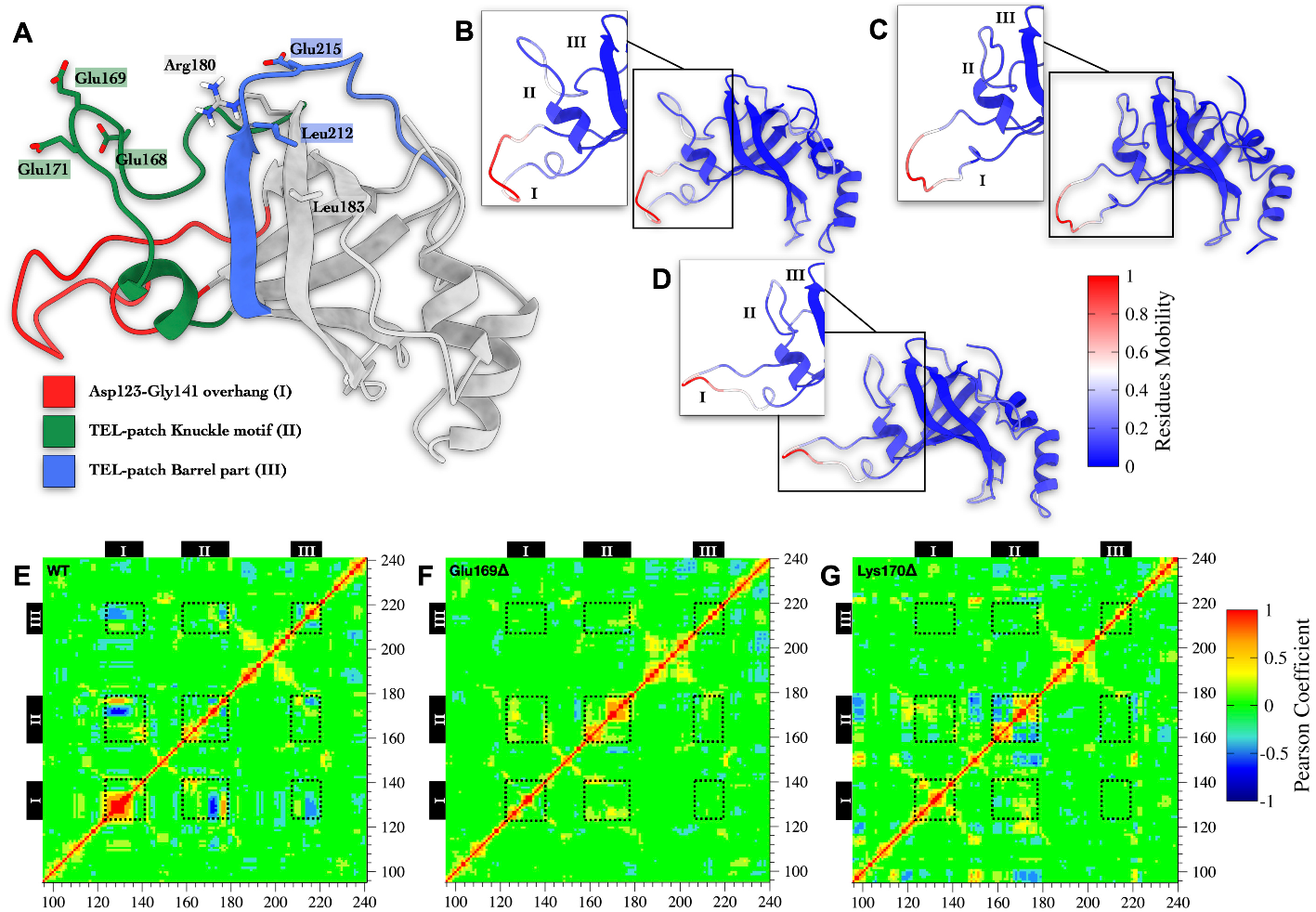
Time series analysis of MD simulations on WT TPP1, Glu169Δ TPP1, and Lys170Δ TPP1 monomers. (A) Tridimensional representation of TPP1’s OB-Domain. (B, C, D) Projection of modules of the first eigenvector computed for WT, Glu169Δ, and Lys170Δ TPP1, respectively. Large scale protein motion with slow timescale is depicted in red. (E, F, G) Pearson coefficients computed between pairs of residues in WT, Glu169Δ, and Lys170Δ TPP1 during the MD calculations, respectively. Values larger than 0.6 are displayed in red (strong correlation), while those smaller than −0.6 are displayed in blue (strong anti-correlation). Residues forming the relevant areas of TPP1 OB-domain are indicated as black dashed line squares.

In Fig. 3B-D, we report the projection of the module of the first eigenvector onto the protein structures for WT, Glu169Δ, and Lys170Δ TPP1, respectively. This represents the slowest and largest conformational change in the protein structure. It is worth noting that a significant motion of **II** only occurs in WT TPP1, while a more rigid structure characterizes the two variants. Similar behavior is also found for **I**, which is endowed with a reduced flexibility in both Glu169Δ and Lys170Δ TPP1. Along with PCA, we performed a cross-correlation analysis (CCA), which allows identifying short- and long-range allosteric effects between residues [26]. In such a way, we could elucidate possible communication networks in the OB-domains. Regarding WT TPP1, protein regions **I**, **II** and **III** show anti-correlated motions - i.e., negative Pearson coefficients - each with the other (Fig. 3E). On the other hand, these signals are lost in the OB-domain variants (Fig. 3F and 3G). This analysis confirms the presence of a concerted motion in the WT between the TEL-patch region and the rest of the protein, while the deletions in the TEL-patch strongly affect the conformational freedom of Glu168-Glu171 in region **II**, as also suggested in a previous study [22]. A complementary result was obtained by calculating the Protein Structure Networks (PSNs) for both WT TPP1 and its variants, using a graph-based approach that assesses the strength of the inter-residue interactions along an MD trajectory [27]. In detail, the PSN of WT TPP1 is characterized by a higher number of edges with respect to the PSN of Glu169Δ TPP1 and Lys170Δ TPP1 (see Fig. S2A-C). Moreover, a distinguishing feature of the variants is the dense communication network between the regions **II** and **III** of the TPP1’s OB-domain, in agreement with the finding that TEL-patch assumes a *closed conformation* in such forms. Overall, the deletion of both Glu169 and Lys170 induces a lower number of intra-protein connections (see Fig. S2D-F), expression of a reduced protein plasticity. Taken together, our findings indicate that the formation of long-lasting interactions between residues at region **II** abolishes the concerted motion observed in WT TPP1, highlighting the impact of even single-point deletions on the entire protein dynamics as in the case of Glu169Δ and Lys170Δ variants.

### 2.3 Effect of TPP1 variants on the formation of WT TPP1-TERT complex

The recently resolved cryo-EM structure of TPP1-TERT heterodimer (PDB ID: 7TRE) prompted us to investigate the impact on the formation of TPP1-TERT binding complex of the TPP1 variants reported in literature to be associated with pathological phenotypes (see Supplementary Tab. S1). This includes the previously introduced Glu169Δ and Lys170Δ variants, plus the Leu95Gln mutant, which is a mutation far from the TEL-patch [28]. The purpose is to provide the molecular basis for a rational understanding of the effect of such variants on the recruitment of TERT by the shelterin protein TPP1. To this end, we performed protein-protein docking calculations between each variant of TPP1 and TERT, and analysed the docking solutions showing the lower RMSD values with respect to the experimental TPP1-TERT structure (PDB ID: 7TRE) [24] (see Methods for details).

As displayed in Fig. 4A, the deletions Lys170Δ and Glu169Δ affect the folding of the α-helix Trp167-Glu171, which is at the binding interface in the TPP1-TERT complex. Consequently, here some crucial inter-protein interactions are lost. In particular, the WT TPP1-TERT complex is characterized by the TPP1’s Glu168-TERT’s Arg774 salt bridge, and by the charge-enforced H-bond between TPP1’s Glu169 and the side-chain of TERT’s Ser134. The deletion of Lys170 inhibits the proper folding of the Trp167-Glu171 sequence, leading to a slightly different TPP1-TERT binding mode (RMSD w.r.t 7TRE ~ 2.6 ^Å^), where TPP1’s Glu168 and TERT’s Arg774 are no longer able to bind each other. The alteration of the α-helix Trp167-Glu171 is even more evident in the case of the Glu169Δ variant, where the H-bond between TPP1’s Lys170 and TERT’s Ser134 is lost (RMSD w.r.t 7TRE ~ 3.1 ^Å^). On the other hand, although the mutation Leu95Gln is not at the TEL-patch, it turns out to affect the correct TPP1-TERT binding mode. In fact, such mutation alters the hydrophobic interactions composed of TPP1’s Leu95 and Leu181, which favour the formation of the salt bridge between TPP1’s Arg92 and TERT’s Asp129 (see Fig. 4B). In particular, the Leu95Gln mutation induces the formation of the intraprotein H-bond between Gln95 and Asp148, leading to a displacement of TPP1’s Arg92 (RMSD w.r.t 7TRE ~ 1.7 Å), with the consequent loss of TPP1’s Arg92 and TERT’s Asp129 interaction at the binding interface. In order to validate our findings, we performed as “blank test” a docking calculation between the wild type forms of TPP1 and TERT, which confirms the experimental binding mode of the complex 7TRE. Overall, our results indicate that all the Leu95Gln, Glu169Δ, and Lys170Δ variants significantly affect the binding mode and binding affinity of TPP1 to TERT (Fig. 4C).

**Fig. 4.**
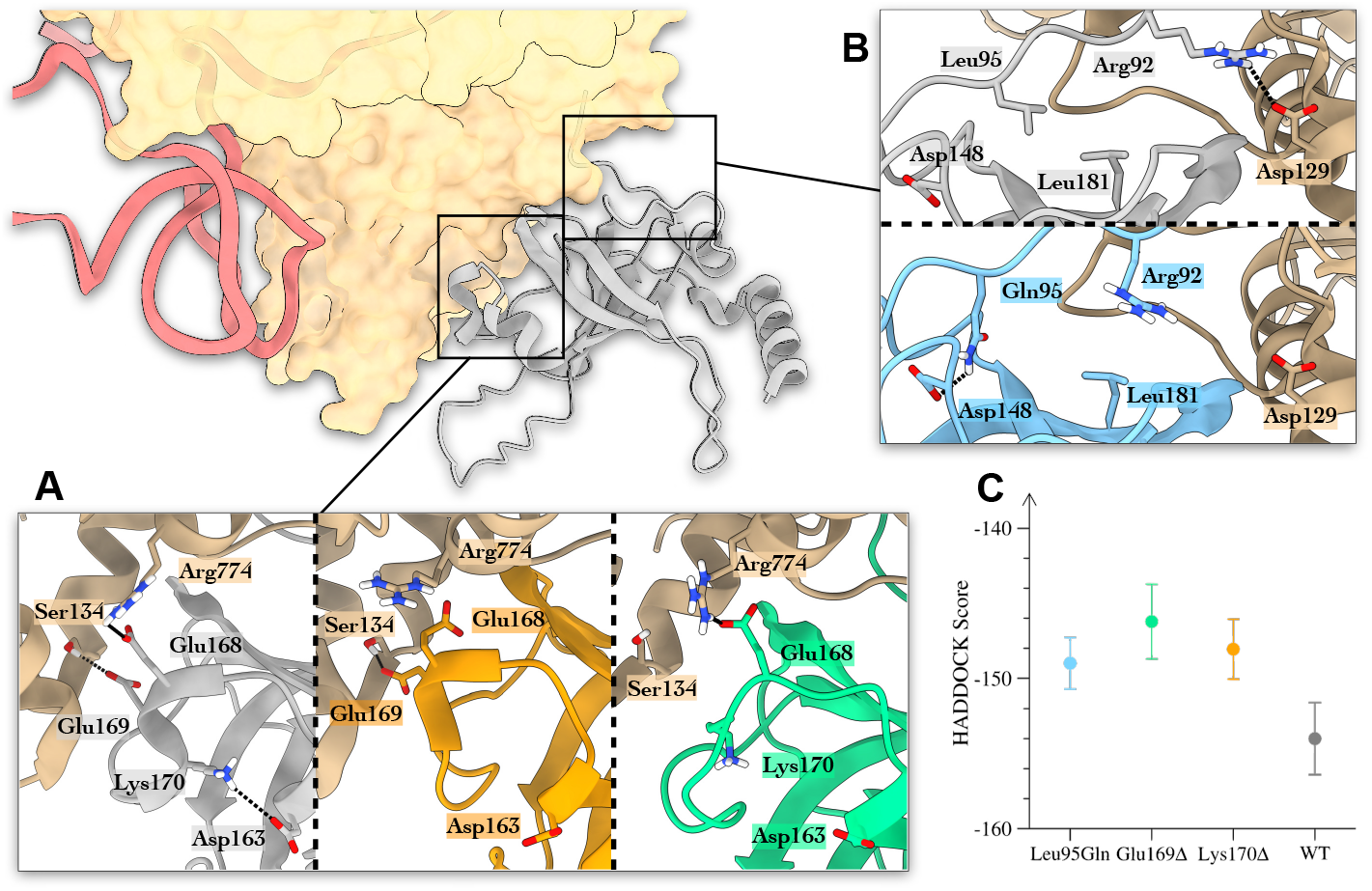
Results of docking calculations performed between TERT and diverse TPP1 mutants. (A) Glu169Δ and Lys170Δ mutants. The interactions hTEN’s Ser134-TPP1’s Glu169 and hTEN’s Arg774-TPP1’s Glu168 are formed thanks to the intraprotein salt bridge between TPP1’s Asp163 and Lys17O (left panel). When such interactions are lost due to Glu169Δ and Lys170Δ deletions, the TPP1-TERT interaction is weakened (central and right panel, respectively). (B) The Leu95Gln mutant. In the wild type TPP1-TERT complex a salt bridge is formed between hTEN’s Asp129 and TPP1’s Arg92, favoured by the hydrophobic stacking between TPP1’s Leu95 and Leu181 (upper panel). The Leu95Gln mutant losses such interaction due to the formation of an interaction between TPP1’s Gln95 and Asp148 that distantiates TPP1’s Arg92 from hTEN’s Asp129 (lower panel). (C) The affinity scores of the best binding modes identified as those with the lower RMSD values with respect to the TPP1-TERT complex reported in PDB ID 7TRE. WT TPP1 is coloured in grey, Leu95Gln TPP1 is coloured in cyan, Glu169Δ TPP1 in orange, and Lys170Δ TPP1 in orange. TERT in coloured in beige and represented through its solvent accessible surface, whereas the TR is shown in magenta.

## 3 Methods

### 3.1 MD simulations on WT TPP1, Glu169Δ TPP1, and Lys170Δ TPP1 monomers

The 3D structure of the TPP1 OB-domain wild type (WT TPP1) was obtained from the protein databank RCSB PDB (PDB ID 2I46) [29]. All the nonprotein species in the crystal structure were removed. The first 5 amino acids (Ser90-Val94) at N-terminal region were deleted since they are unstructured in the original pdb. The 3D structure of Glu169Δ TPP1 was built with the MODELLER suite [30], removing the Glu169 from the original sequence. The same approach was followed to produce the initial structure of Lys170Δ TPP1. In all systems, the N-terminal and C-terminal were capped with an acetyl and a methyl-amino protecting group, respectively. Then, the proteins were solvated using the TIP3P water model, and a salinity of 0.15 M NaCl. The Amber ff99sb-ildn force field was employed [31] with the MD engine GRO-MACS 2016.5 [32]. Each system underwent a thermalisation cycle increasing the temperature from 100 K until 300 K with steps of 50 K, decreasing at each step the restraints applied on the system’s heavy atoms. This setting permits solving possible steric clashes between atoms, however retaining the overall tertiary structure of the proteins. All systems experienced the following protocol: 200 ps of NVT simulation followed by 200 ps of NPT simulation for each step of temperature increase. The V-rescale thermostat was employed in the thermalisation phase, while the Langevin thermostat was used during the production runs. Periodic boundary conditions have been applied and the particle-mesh-Ewald (PME) method was used to treat long-range electrostatic interactions [33]. For short range atom-atom interactions a cut-off distance of 1.0 nm was applied. The pressure was maintained at the reference value of 1 bar using the Parrinello-Rahman barostat [34].

### 3.2 Cluster analysis

Cluster analyses on the MD trajectories were performed using GROMACS’s *“gmx cluster*” routine. The cluster families of WT TPP1, Glu169Δ TPP1, and Lys170Δ TPP1 were obtained by computing the RMSD for the Cα atoms of the TPP1’s residues from Ser165 to Leu184, from Ser210 to Glu215, and the heavy atoms of the side chains belonging to the TEL-patch’s amino acids. The RMSD cut-off was set at 1.8 ^Å^.

### 3.3 Protein-protein contacts definition

In order to assess both TPP1’s intraprotein interactions and the contacts formed by TPP1 and hTEN at the dimer interface, we computed the frequency of occurrence for each contact, represented as histograms using the *“PLOT NA”* routine of Drug Discovery Tool (DDT) [35]. We set 4.0 ^Å^ as the cutoff distance between two interacting residues.

### 3.4 Cross-correlation analysis

Cross-correlation analysis (or Pearson-correlation coefficient analysis) was used to assess the correlated motions between pairs of residues in the monomer and dimer systems. An *in-house* algorithm was employed to calculate the Pearson coefficients matrix for each couple of residues according to the following formula:

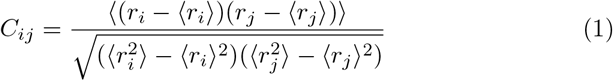

where *r_i_* and *r_j_* are the position vectors of the Cα atoms in residues *i* and *j*, respectively. The angle brackets denote time averages computed along the simulations. The final *C_ij_* value ranges from −1.0 to 1.0. The higher the *C_ij_* value, the stronger is the linear correlation of the motion of that pair of residues.

### 3.5 Graph-based network analysis

The Protein Structure Network (PSN) approach was employed to assess the effect of the Glu169Δ and Lys170Δ mutations on TPP1’s network connectivity. Network parameters such as hubs, communities, and structural communication analyses were obtained by using the WebPSN 2.0 web-server [36–38]. The methodology builds the Protein Structure Graph (PSG) based on the interaction strength of two connected nodes:

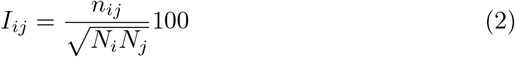

where interaction percentage (I_ij_) of nodes i and j represents the number of pairs of side-chain atoms within a given cut-off value (4.5 ^Å^), while *N_i_* and *N_j_* are normalization factors. The interaction strength (represented as a percentage) between residues *i* and *j* (*I_ij_*) is calculated for all node pairs. If *I_ij_* is more than the minimum interaction strength cutoff (*I_min_*) among the residue pairs, then is considered to be interacting and hence represented as a connection in the PSG. The graphs shown in Fig. S1 and S14 have been generated using the *“networkx”* library of python3 [39].

### 3.6 Docking on TPP1-TERT dimers

The starting conformations of Leu95Gln TPP1, Glu169Δ TPP1, and Lys170Δ TPP1 were generated through the MODELLER suite, employing TPP1’s 7TRE as the initial template [24]. The 3D coordinates of the complex formed by Leu95Gln TPP1 with TERT, Glu169Δ TPP1 with TERT, and Lys170Δ TPP1 with TERT were obtained by using the molecular docking HADDOCK 2.4 webserver [40]. For the docking calculations, we indicated TPP1’s Arg92, Glu168, Leu183, Leu212, Pro213, Glu215, and TERT’s Lys78, Asp129, Arg132, Leu766, Leu769, Tyr772, Arg774 as points of contact (i.e., active residues), in line with the recently released structure of the TPP1-TERT complex. The best binding poses were selected as those having the lowest RMSD with respect to 7TRE.

## 4 Conclusion

The molecular binding interaction between the Shelterin protein TPP1 and telomerase is a key player in telomere maintenance mechanism and genome protection. The recently resolved cryo-EM structure of TPP1-telomerase complex (PDB ID 7TRE) [24] has represented a breakthrough in the field, providing the structural basis for understanding the binding interaction between the two proteins. However, experimental structures do not provide information on protein functional dynamics that are relevant for a comprehensive understanding of the molecular recognition process and for rationalizing the effects of pathological variants of the complex. Examples are the TPP1 single-point deletions Glu169Δ and Lys170Δ that induce telomere shortening and accelerated ageing in Hoyeraal-Hreidarsson syndrome. Atomistic simulations are apt to this scope, being able to complement the experimental structures - obtained by cryo-EM, X-ray or NMR - with the missing conformations for a comprehensive understanding of the protein functional mechanism. Here, we show by combining microsecond-scale MD calculations, multivariate analysis, cryo-EM, and mutagenesis data that the TPP1’s OB-domain is endowed with significant conformational plasticity, regulated by allosteric communications between three regions **I**, **II**, **III**, distant in the protein structure (Asp123-Gly141 overhang, TEL-patch Knuckle motif and TEL-patch barrel part, respectively). Such communication network stabilises the TEL-patch region in *a closed* and a *open conformation*, the latter functional for the TEL-patch binding to TERT. Interestingly, such network is lost in the Glu169Δ, Lys170Δ and Leu95Gln pathological variants of TPP1, where the conformational rigidity of the *“Knuckle*” motif in the TEL-patch (region **II**) does not permit its correct folding in an α-helix at the binding interface. As a result, the known landmark interaction formed between TPP1’s Glu215 and TERT’s Lys78 is lost, affecting the correct binding with hTEN that might reduce telomere processivity and cause accelerated ageing. The atomistic structures of wild type and mutated TPP1 in complex with telomerase - released as PDB files in SI - pave the way to rational drug discovery studies aimed at restoring the correct heterodimer formation. One could imagine designing allosteric ligands that target the three TPP1 regions and stabilise the open state of the TPP1 OB-domain competent for the recruitment of hTEN. Furthermore, the heterodimer complex formed by TPP1 and hTEN represents an attractive molecular target to develop anticancer agents. In this case, TEL-patch targeting ligands might inhibit the TPP1-telomerase binding and induce cancer cell senescence. Using such an approach, we have recently discovered the first TPP1 ligands with anticancer activity whose pharmacological characterization is underway. In conclusion, our results endorse atomistic simulations as a valuable tool to disclose TPP1 functional dynamics as shown in other systems [41–44]. The TPP1-telomerase structures here presented represent an attractive pharmacological target for drug design of innovative therapeutics in cancer and telomeropathies like the Hoyeraal–Hreidarsson syndrome.

## Supporting information

TPP1_structures

Video_TPP1

## Supplementary information

Atomic coordinates of the TPP1 structures W1, W2, W3, ΔE1, ΔE2, ΔK1, and ΔK2 are available in the Supplementary Materials in the “TPP1_structures.zip” file. A video detailing the effect of Lys170Δ on TPP1 functional dynamics is available at the following link and as Supplementary Materials.

## Acknowledgments

The authors thank Robert L. Jernigan and Daniele Di Marino for useful discussions. We acknowledge the support from the European Research Council (ERC Consolidator Grant “CoMMBi”), the Swiss National Science Foundation (Project No. 200021_163281), the Italian MIUR-PRIN 2017 (2017FJZZRC), the Swiss National Supercomputing Centre (CSCS) [project ID s1116], and the Trés Grand Centre de Calcul (TGCC) [project ID ra4681].

