## Supplementary Information for "Conformational plasticity and allosteric communication networks govern Shelterin protein TPP1 binding to human telomerase"

**SUPPLEMENTARY MATERIAL**

The data hereby reported are divided in the following sections:

- Plot of the intra-protein interactions in WT TPP1, Glu169Δ TPP1, and Lys170Δ TPP1 monomers
- Protein structure network of the WT TPP1, Glu169Δ TPP1, and Lys170Δ TPP1 monomers
- List of TPP1 mutations

**Intra-protein interactions in WT TPP1, Glu169Δ TPP1, and Lys170Δ TPP1 monomers**

In this section, we displayed our analysis of the time evolution of the intra-protein contacts ruling the conformational plasticity of the TEL-patch, i.e. Glu171-Arg180, Glu169-Arg180, Glu168-Arg180, and Asp166-Ser210. As shown in Fig. S1, the MD simulations of Glu169Δ TPP1 and Lys170Δ TPP1 are marked by the establishment of lost-lasting interactions along the whole trajectories.

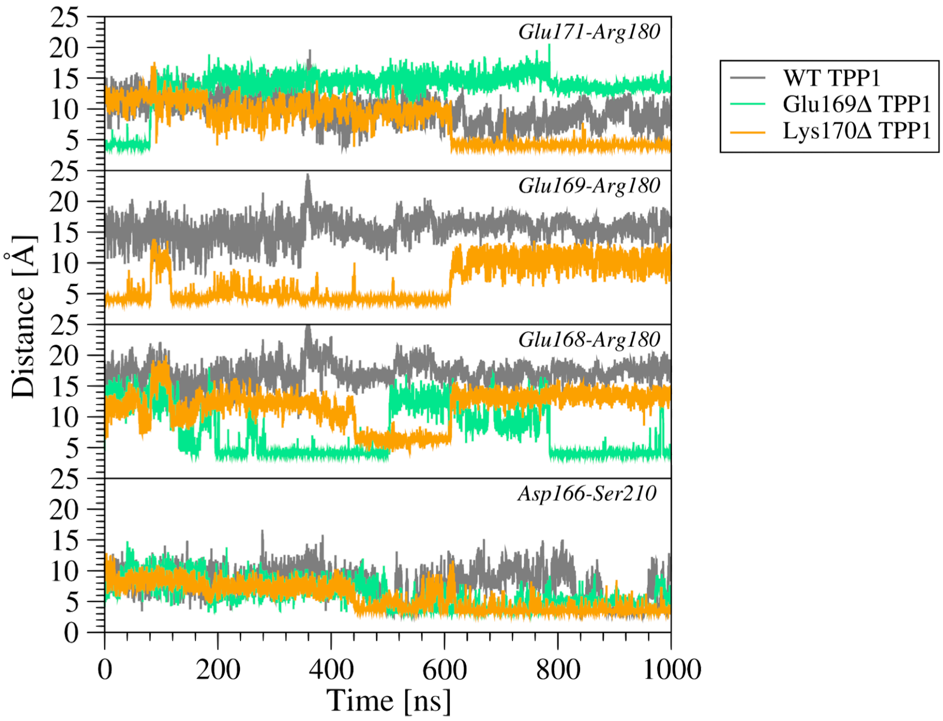


**Fig. S1:** Plot reporting the intra-protein distances ruling the conformational plasticity of the TEL-patch, i,e, Glu171-Arg180, Glu169-Arg180, Glu168-Arg180, and Asp166-Ser210. The interactions occurring during the WT TPP1 simulation are coloured in grey, while the ones established in Glu169Δ TPP1 and Lys170Δ TPP1 are coloured in green and orange, respectively.

**Protein structure network of the WT TPP1 and Lys170Δ TPP1 monomers**

In this section, we reported our analysis of the WT TPP1's, Glu169Δ TPP1's, and Lys170Δ TPP1's Protein Structure Network (PSN). In PSN, the residues are considered as nodes in a graph, interconnected by edges with attributed weights that are computed based on the non-covalent atomistic contacts established between nodes. In such a way, it is possible to compare graphs representing the same protein at different conditions. We have exploited this possibility by resolving the exclusive paths of TPP1 in the different systems, that is the identification of inter-residue interactions in TPP1 that are only present in the WT and in the Glu169Δ/Lys170Δ variants. As shown in Fig. S2 reported below, the PSN analysis agrees with the results obtained through the calculation of Pearson coefficients reported in Fig. 3, confirming the loss of most of the inter-residue interactions between the TEL patch and the Asp123-Gly141 overhang in the TPP1 mutants with respect to the WT.

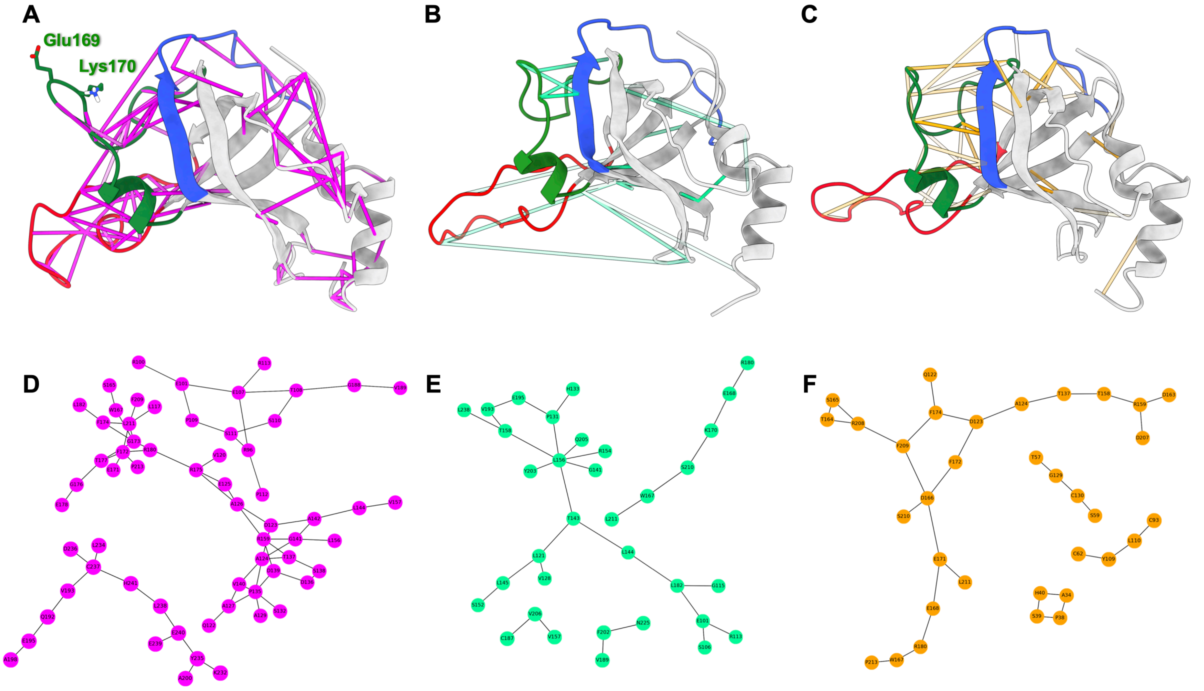


**Fig. S2:** Graph-based network analysis of the monomeric structure of WT TPP1, Glu169Δ TPP1, and Lys170Δ TPP1 during MD simulations. (**A**-**C**) The communications between the residues are projected upon the 3D protein structure of WT TPP1 (**A**), Glu169Δ TPP1 (**B**) and Lys170Δ TPP1 (**C**). (**D**-**F**) Representation of the graphs corresponding to WT TPP1, Glu169Δ TPP1, and Lys170Δ TPP1, respectively. The three relevant moieties of the TPP1's OB domain ((**I**) Asp123-Gly141 overhand motif; (**II**) TEL-patch Knuckle motif; (**III**) TEL-patch Barrel part) are coloured following the color scheme reported in Fig. 3 of the original manuscript. The graph edges of WT TPP1 are coloured in magenta, the edges for Glu169Δ TPP1 are coloured in grees, while the edges for Lys170Δ TPP1 are in orange. Transparency is proportional to the weight of the edge, i.e. solid edges correspond to highly frequent connections. The graphs reported in panels **D**, **E**, and **F** are generated to best represent the connection between nodes and are unrelated to the protein 3D structure.

### List of TPP1 mutations

Table S1 reports the known TPP1 mutations at their binding interface with TERT with the corresponding phenotypes and references.

| TPP1's mutations | Phenotype | Reference |
|---|---|---|
| Leu95Gln | Severe telomeres shortening | [1] |
| Glu169Δ | Reduced telomeres length-HHS | [2-3] |
| Lys170Δ | Reduced telomeres length-HHS | [4-6] |

**Tab. S1**: Tables reporting the TPP1 mutations involving residues the interface of the WT TPP1-hTEN dimer interface (as depicted in PDB ID: 7TRE). The impact of all these mutations on the heterodimer stability was verified through docking calculations. For each mutation, we indicated both the corresponding experimental phenotype and the related references.
